# Characterizing functional consequences of DNA copy number alterations in breast and ovarian tumors by spaceMap

**DOI:** 10.1101/248229

**Authors:** Christopher J. Conley, Umut Ozbek, Pei Wang, Jie Peng

## Abstract

**Motivation:** We propose a novel conditional graphical model — `spaceMap` — to construct gene regulatory networks from multiple types of high dimensional omic profiles. A motivating application is to characterize the perturbation of DNA copy number alterations (CNA) on downstream protein levels in tumors. Through a penalized multivariate regression framework, `spaceMap` jointly models high dimensional protein levels as responses and high dimensional CNA as predictors. In this setup, `spaceMap` infers an undirected network among proteins together with a directed network encoding how CNA perturb the protein network. `spaceMap` can be applied to learn other types of regulatory relationships from high dimensional molecular profiles, especially those exhibiting hub structures.

**Results:** Simulation studies show `spaceMap` has greater power in detecting regulatory relationships over competing methods. Additionally, `spaceMap` includes a network analysis toolkit for biological interpretation of inferred networks. We applied `spaceMap` to the CNA, gene expression and proteomics data sets from CPTAC-TCGA breast (n=77) and ovarian (n=174) cancer studies. Each cancer exhibited disruption of ‘ion transmembrane transport’ and ‘regulation from RNA polymerase II promoter’ by CNA events unique to each cancer. Moreover, using protein levels as a response yields a more functionally-enriched network than using RNA expressions in both cancer types. The network results also help to pinpoint crucial cancer genes and provide insights on the functional consequences of important CNA in breast and ovarian cancers.

**Availability:** The R package `spaceMap` — including vignettes and documentation — is hosted at https://topherconley.github.io/spacemap

## 1. Introduction

Relative to high-throughput genomic assays, only recently has quantitative mass-spectrometry-based proteomics become available for large-scale studies. The collaboration between the Clinical Proteomic Tumor Analysis Consortium (CPTAC) (Paulovich et al., 2010; Ellis et al., 2013; Zhang et al., 2016) and The Cancer Genome Atlas (TCGA) is among the first to produce large-sample cancer studies integrating deep proteomic and genomic quantitative profiling. Naturally this proteogenomic combination enables the study of how and to what extent genetic alterations impacted protein levels in cancer. Specifically, the CPTAC Breast/Ovarian Cancer Proteogenomics Landscape Study (BCPLS/OCPLS) identified proteomic signaling consequences of DNA copy number alterations (CNA) based on 77 and 174 high-quality breast and ovarian cancer samples, respectively (Mertins et al., 2016; Zhang et al., 2016). Identifying which CNAs impact downstream protein activities and how they do so can lead to better understanding of disease etiology and discovery of new biomarkers as well as drug targets (Akavia et al., 2010; Greenman et al., 2007).

However, the full extent of biomedical information in multiple-omic studies like BCPLS/OCPLS cannot be realized without effective integrative analysis tools. These tools must characterize interactions among different biological components operating in different cellular contexts. For the BC-PLS/OCPLS data and other proteogenomic data like it, this requires examining relationships between a large number of protein levels in one context and CNA events in another context. Studying a pair of features at a time may not be sufficient, as cancer is overwhelmingly complex; molecules from many parallel signal transduction pathways are affected by the disease, and their activities appear to be controlled by multiple factors. Therefore, we need to jointly consider all players in the system. Recent advances in graphical models for high-dimensional data provide a powerful “hammer” for this “nail” (Meinshausen and Bühlmann, 2006; Yuan and Lin, 2006; Friedman et al., 2008; Peng et al., 2009).

Graphical models infer interactions among features (e.g., genes/proteins) based on their dependency structure, for it is believed that strong interactions often result in significant dependencies. Compared to approaches using pairwise correlation to characterize and infer interacting relationships (e.g. (Butte et al., 2000)), graphical models learn more direct interactions through investigating *conditional dependencies*. Many methods have been proposed in the past decade to infer *genetic regulatory networks* (*GRNs*) based on high-throughput molecular profiling through graphical models (Li et al., 2013; Friedman et al., 2008; Peng et al., 2009; Wang et al., 2011; Cheng et al., 2014; Danaher et al., 2014; Schäfer and Strimmer, 2004). However, when applying graphical models for the proteogenomic integrative analysis, we encounter new challenges that are not fully addressed by existing methods. First, integrative analysis necessarily involves multi-layer biological components, but we may only be interested in a subset of all possible interactions. For example, we may be interested in how CNAs regulate protein levels, but not the dependency structure among CNAs. Second, although many data types may be reasonably modeled by Normal distributions, some data types, such as DNA mutation and SNP, can not be modeled by Gaussianity.

To bridge these gaps we propose a conditional graphical (CG) model, `spaceMap`, which learns the conditional dependencies between two types of nodes through a penalized multivariate regression framework. Specifically, `spaceMap` infers an undirected graph among response variables (e.g., protein levels) in tandem with a directed graph encoding perturbations from predictor variables (e.g., CNAs) on the response network. In addition, we use cross-validation and a model aggregation technique called Boot.Vote to improve reproducibility. Moreover, we develop a network analysis toolkit to facilitate biological interpretation. These lead to an integrative -omics analysis pipeline illustrated in Figure 1.

**Figure 1:**
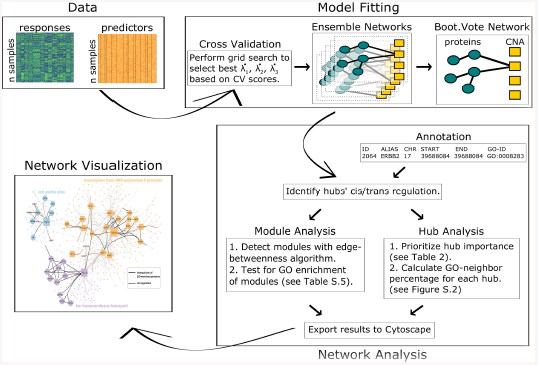
spaceMap integrative analysis pipeline. Predictors (e.g., CNA) and responses (e.g., protein abundance) data are inputs to the model fitting stage, where the model is tuned by cross validation and aggregated across 1000 bootstrap ensemble networks through the Boot.Vote procedure. The Boot.Vote network is input to the network analysis stage, where biological function is layered onto the network. Finally the network is visualized with Cytoscape.

Peng et al. (2010) proposed the `remMap` model for integrative analysis of gene expressions and CNA. Specifically, `remMap` utilized a penalized multivariate regression model and introduced a so called MAP penalty through combining both *l*_1_ and column-wise-*l*_2_ norm of the coefficient matrix to encourage the selection of *master* predictors that have influence on many responses. However, unlike `spaceMap`, `remMap` does not reveal how the expressions interact with each other in the presence of perturbations from CNAs.

Another straight forward approach to handle two types of data is to fit one graphical model without distinguishing the two types of nodes and then subset only those interactions of interest. For example, a model like space (Peng et al., 2009) can be used to infer an undirected network where CNAs and protein levels are not explicitly distinguished during model fitting. The inferred conditional dependencies amongst CNAs can then be ignored, while CNA–protein and protein–protein interactions could be retained for further investigation. However, since there are often thousands of CNA regions, such an approach suffers from inefficiency by wasting many degrees of freedom in estimating interactions among CNAs that we are not interested in. On the contrary, `spaceMap` aims at learning the conditional dependencies among a set of response nodes (e.g., protein levels) and the perturbations from a set of predictor nodes (e.g., CNAs). By conditioning on the predictor nodes, these are free to have any distribution and the interactions among the predictors need not be modeled and estimated. Such an approach is expected to exhibit gains in statistical power and computational efficiency.

Recently, Zhang and Kim (2014) proposed a model called `scggm` to fit conditional Gaussian graphical models through an *l*_1_ penalized conditional log-likelihood. In several genetical genomics simulations, `scggm` showed higher precision-recall curves (i.e. accuracy) in learning the network structure than competing methods including `MRCE` (Rothman et al., 2010) and `graphical lasso` (Friedman et al., 2008). Although `spaceMap` and `scggm` target the same type of response-predictor and response-response interactions, they are very different in terms of modeling approaches. `spaceMap` uses a regression-based approach through pseudo-likelihood approximations with a penalized least squares criterion, while `scggm` is based on penalized conditional likelihood. Peng et al. (2009) conducted a comprehensive comparison between regression-based and likelihood-based graphical models and found that regression-based methods often perform better in the presence of hubs and are more robust to violations of distributional assumption. Moreover, `spaceMap` adopts the MAP penalty from `remMap` (Peng et al., 2010) for the response-predictor interactions, making it more powerful in detecting master predictors. In a simulation study involving networks with hub nodes, `spaceMap` is shown to outperform both `space` and `scggm` in terms of edge and hub detection.

The rest of the paper is organized as follows. In Section 2, we first present simulation results to demonstrate the performance of `spaceMap` and then apply `spaceMap` to the BCPLS data and OCPLS data to learn the (CNA–protein, CNA-RNA, and protein–protein) regulatory networks. We summarize the conclusions in Section 3. In Section 4, we describe the `spaceMap` model, simulation settings, BCPLS/OCPLS applications, and a network analysis toolkit. Additional details are given in the Supplementary Material.

## 2. Results

### 2.1. Simulation

Figure 2 shows the results of the three methods, namely `spaceMap, space` and `scggm`, under the `hub-net` simulation (see Section 4.2). `spaceMap` has the highest MCC and the highest power across all edge types while maintaining a low FDR. Particularly, `spaceMap` has considerably more power in CNA–protein edge detection than the other two methods. `space` is least powerful in CNA–protein edge detection, though it also has the lowest FDR. The differences in power and MCC are significant, whereas those in FDR are not always significant. See Supplementary Table S.1 for more details.

**Figure 2:**
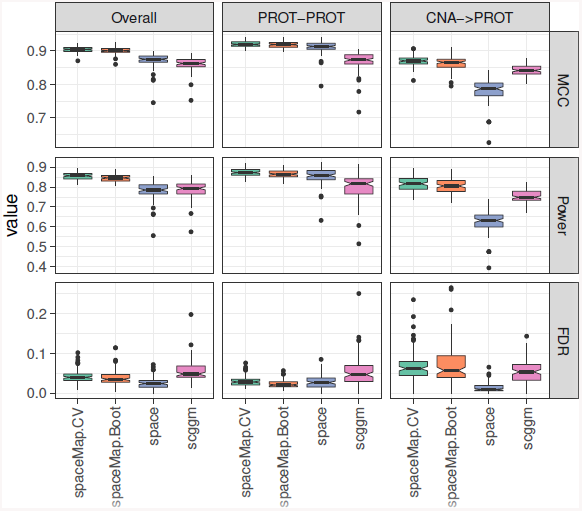
hub-net simulation. Edge-detection performance summarized by MCC, power, and FDR, across 100 replicates. The overall performance is further decomposed into response subnetwork PROT–PROT and the predictor→response subnetwork CNA→PROT. `spaceMap`.CV, space and scggm are learned under CV.Vote and `spaceMap`.boot is learned under Boot.Vote. All tuning parameters are chosen by 10-fold CV.

Moreover, all methods have 100% power in identifying all 15 CNA-hubs. Note that, the CNA-hub power is defined as the power to detect at least one CNA-protein interaction. Since each hub has a fair number of such interactions, this power is expected to be high for all reasonable methods. On the other hand, `spaceMap` has the lowest CNA-hub detection FDR (0.67%) and `scggm` has the highest CNA-hub detection FDR (12.6%).

The performance of the three methods under the `power-net` simulation (see Section 4.2) is shown in Figure S.1 with numerical summaries given in Table S.2 of the Supplementary Material. `spaceMap` achieves the highest MCC score across all edge types followed with a close second by space, and a distant third by `scggm`. `spaceMap` dominates space and `scggm` in CNA–protein edge detection power at the cost of slight FDR inflation. `scggm` has the highest power in protein–protein edge detection, however with extremely high FDR. All methods exhibit (near) 100% power in CNA-hub detection. Remarkably, `spaceMap` has an CNA-hub FDR of 0, while the other two methods have CNA-hub FDR above 20%. In summary, `spaceMap` is able to find the true source of perturbation to the response network. On the other hand, the other two methods exhibit a tendency to report false CNA-hubs.

We also apply the Boot.Vote procedure (with *B* = 1000 bootstrap resam-ples) described in Section 4.1 to `spaceMap` to further reduce FDR. `Boot.Vote` under the `hub-net` simulation leads to very similar results as `CV.Vote`, probably due to the already low FDR in this setting. On the other hand, `Boot.Vote` under the `power-net` simulation leads to reduced FDR compared to `CV.Vote.` In short, `Boot.Vote` is an effective procedure in (further) reducing FDR. It is especially useful for real data applications, which often suffer from low signal-to-noise ratio and complicated noise structure and consequently high FDR and low reproducibility of the fitted networks.

### 2.2. BCPLS application

#### 2.2.1. prot-net

We first focus on protein levels because protein activities are expected to have a more direct impact on cell phenotypes than RNA. The goal is to identify major CNA events disrupting biological pathways at the protein level while accounting for the conditional dependency structure among proteins themselves. To this end `spaceMap` learns a network from CNA and protein levels — hereafter called `prot-net` — built under the 10-fold CV-selected tuning parameters and the `Boot.Vote` (*B* = 1000) aggregation process.

`prot-net` has 585 CNA–protein edges, 954 protein–protein edges and 11 CNA-hubs (Table 1). The top three ranked CNA-hubs are listed in Table 2 and the complete list is provided in Table S.4. Network analysis reveals 10 modules of size 15 or more. GO terms enriched in each module are listed in Supplementary Table S.5. Three of the 10 modules contain at least one CNA-hub that directly perturbs at least five proteins within the module. Figure 3 illustrates the topology of these three modules.

**Figure 3:**
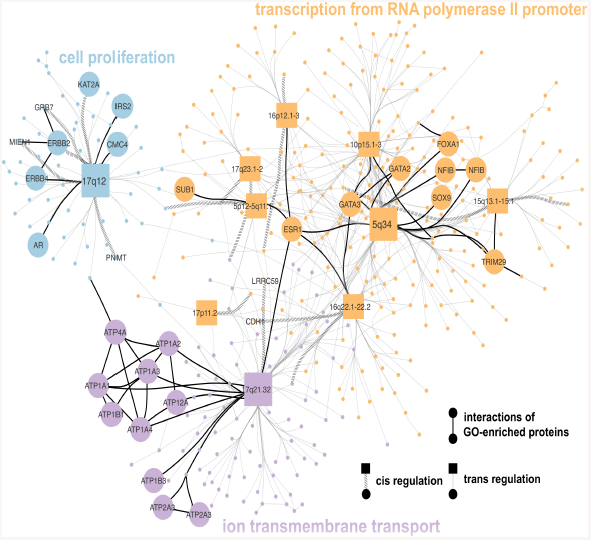
BCPLS application. Three GO-enriched modules from `spaceMap` `prot-net`. Large circles denote proteins belonging to enriched GO terms: cell proliferation (blue), transcription from RNA polymerase II promoter (orange) and ion transmembrane transport (purple). Rectangles denote CNA-hubs.

**Table 1:** Summary statistics of `prot-net` and RNA-net by `spaceMap`. For CNA-hubs and GO-neighbor percentage, statistics are computed on CNA-hubs with at least degree 10 (prior to manually merging a few highly correlated CNA-hubs).

| Statistic | Breast cancer |  | Ovarian cancer |  |
| --- | --- | --- | --- | --- |
|  | prot-net | RNA-net | prot-net | RNA-net |
| # CNA nodes | 1662 | 1662 | 1349 | 1349 |
| # expr nodes | 1595 | 1657 | 2097 | 2236 |
| # expr-expr edges | 954 | 1010 | 1657 | 1735 |
| # CNA-expr edges | 585 | 622 | 1284 | 1231 |
| # CNA-hubs (median size) | 11 (36) | 9 (55) | 6 (10) | 44 (17.5) |
| median GO-neighbor % | 71.43 | 57.90 | 50.0 | 52.77 |

**Table 2:** BCPLS/OCPLS applications. Top three CNA-hubs for `prot-net` and RNA-net of breast cancer and ovarian cancer, respectively. Within each CNA-hub, we report the number of cis- and trans- edges and the potential number of cis regulations (i.e., the number of protein/RNA nodes in cis with this CNA-hub from the entire network). We also list which genes are found to be in cis with the CNA-hub.

| Cytoband | # cis/<br># trans | Potential<br># cis | cis-<br>proteins |
| --- | --- | --- | --- |
|  | Breast cancer | prot-net |  |
| 17q21.32 (46-46 Mb) | 1 / 98 | 14 | LRRC59 |
| 5q34 (160-170 Mb) | 0 / 129 | 0 | – |
| 17q12 (38-38 Mb) | 5 / 47 | 34 | ERBB2, GRB7, MIEN1<br>PNMT, KAT2A |
|  | Breast cancer | RNA-net |  |
| 5q34 (160-170 Mb) | 0 / 189 | 4 | – |
| 10p15.1-15.3 (0.42-4 Mb) | 0 / 55 | 7 | – |
| 17q12 (38-38 Mb) | 4 / 36 | 24 | ERBB2, GRB7, PNMT, TCAP |
|  | Ovarian cancer | prot-net |  |
| 22q13.1-.31 (39-46 Mb) | 5/5 | 26 | TSPO, ACO2, TTLL12,<br>CYB5R3, ARFGAP3, |
| 20q11.22-.23 (32-36 Mb) | 3/16 | 20 | NFS1, RPRD1B, CPNE1 |
| 8q24.23-24.3 (140-150 Mb) | 11/3 | 16 | C8orf82, GSDMD, CYC1,<br>TSTA3,KHDRBS3, THEM6,<br>OPLAH, PYCR3, EPPK1,<br>MROH1, NAPRT |
|  | Ovarian cancer | RNA-net |  |
| 22q13.1-13.31 (39-46 Mb) | 3 / 103 | 11 | KDELR3, APOBEC3B, APOL6 |
| 20q11.22-11.23 (32-36 Mb) | 1 / 103 | 3 | ID1 |
| 8q22.1-23.3 (98-120 Mb) | 3 / 46 | 11 | NCALD, TNFRSF11B, TRPS1 |

One of the three modules contains a CNA-hub on 17q12 (Figure 3 upper-left), which cis-regulates multiple genes in this region including ERBB2, GRB7, and PNMT. ERBB2, the epidermal growth factor receptor 2, is a well-known breast cancer oncogene for Her2-subtype of breast cancer. Activities of Her2, the protein coded by ERBB2, influence multiple pathways regulating cell growth, survival, migration and proliferation that have a key role in cancer development. Her2 has been used as a drug target in current clinical treatment of breast cancer patients (Wolff et al., 2013; Sahlberg et al., 2013). Recent research studies also suggest that expression of other genes in the 17q12 amplicon, such as GRB7 and PNMT, may function together with ERBB2 to sustain the growth of breast cancer cells (Sahlberg et al., 2013). Thus, identifying CNA in 17q12 as a hub in CNA–protein regulatory network suggest that `spaceMap` is able to reveal known important regulations underlying the disease system.

In addition to 17q12, `spaceMap` also identified another prominent CNA regulatory hub on the same chromosome in 17q21.32 (Figure 3, bottom). Loss of 17q21.32 has been widely reported in breast cancer studies. For example, in a recent paper of invasive ductal breast cancer study, loss in 17q21.32 were observed at a very high frequency (80%), suggesting potential tumor suppressor genes harbored in this region (Dimova, 2015). However, it remains unclear what functional consequences these deletions may lead to in tumor cells. To address this question, we investigated the subset of proteins regulated by the CNA-hub on 17q21.32. We identify a tightly linked group of 10 proteins from the family of P-type cation transport ATPases, which are integral membrane proteins responsible for establishing and maintaining the electrochemical gradients of Na and K ions across the plasma membrane. Increasing evidence suggests that ion channels and pumps play important roles in cell proliferation, migration, apoptosis and differentiation, and therefore is involved in aberrant tumor growth and tumor cell migration (Li and Langhans, 2015). For example, Na, K-ATPase proteins are associated with various signaling molecules, including Src, phosphoinositide 3-kinase (PI3K), and EGFR thereby activating a number of intracellular signaling pathways, including MAPK and Akt signaling, to modulate cell polarity, cell growth, cell motility and gene expression (Haas et al., 2002; Barwe et al., 2005). In addition, it has been hypothesized that targeting overexpressed Na(+)/K(+)-ATPase alpha subunits might open a new era in anticancer therapy and bring the concept of personalized medicine from aspiration to reality (Mijatovic et al., 2008). Our network analysis result further suggests that high frequency deletion of 17q21.32 might serve as an upstream regulating event for Na, K-ATPase proteins in breast cancer cells. Further study of this region could lead to new discoveries of tumor suppressor genes controlling ion channels and pumps (Litan and Langhans, 2015). Additionally, 17q21.32 acts in cis on LRRC59, which has been shown to modulate cell motility, aid in EMT, and is a necessary factor for oncogene FGF1 to enter the nucleus for regulatory activity (Maurizio et al., 2016).

Loss of 5q is a common feature of basal-like breast tumors (Cancer Genome Atlas Network et al., 2012). In the network inferred by `spaceMap`, a CNA of a region on 5q34 is found to influence the largest number of proteins (Figure 3, upper-right). This observation is consistent with previously reported result based on pairwise correlation analysis on the same data set (Mertins et al., 2016). Applying GO enrichment analysis to the network module containing the 5q34 CNA-hub reveals the biological process of “regulation of transcription from RNA polymerase II promoter” is significantly enriched, including the well-known breast cancer oncogenes ESR1, GATA3, FOXA1 and many others. No doubt that gene transcription mediated by RNA polymerase II (pol-II) is a key step in gene expression, as the dynamics of pol-II moving along the transcribed region influence the rate and timing of gene expression. Specifically, in breast cancer cell lines, it has been confirmed that the predominant genomic outcome of estrogen signaling is the post-recruitment regulation of pol-II activity at target gene promoters, likely through specific changes in pol-II phosphorylation (Kininis et al., 2009). Another recent work also demonstrate that pol-II regulation is impacted during activation of genes involved in the epithelial to mesenchymal transition (EMT), which when activated in cancer cells can lead to metastasis (Samarakkody et al., 2015). These findings together with our results from `spaceMap` analysis imply that DNA copy number alterations of 5q34 plays important role in breast tumor initiation and progression.

Mertins et al. (2016) reported 6 trans- association hubs at chromosomal arm level, namely, 5q, 10p, 12, 16q, 17q, and 22q. `spaceMap` identified CNA-hubs in all six arms except for 22q. On the other hand, `spaceMap` identified additional tran-hubs in 15q13.1-15.1, 11q13.5-14.1, 12q21.1. These details are provided in Table S.14. `spaceMap` further pinpointed new cis-regulation between the CNA-hub in 17q12 and MIEN1 as well as KAT2A. Migration and invasion enhancer 1 (MIEN1) is an important regulator of cell migration and invasion. In a recently reported study, MIEN1 is found to drive breast tumor cell migration by regulating cytoskeletal-focal adhesion dynamics, and targeting MIEN1 is suggested to be a promising means to prevent breast tumor metastasis (Kpetemey et al., 2016). KAT2A, also known as histone acetyltransferase GCN5, is reported to play an essential role in the HBXIP-enhanced migration of breast cancer cells by wound healing assay, and thus is also an important player in tumor metastasis of breast cancer (Li et al., 2015).

To have a more direct comparison with the marginal correlation based approach, we built a network (referred to as the marginal network) using the same set of nodes as in `prot-net` (Table 1) whose edges were determined by significant Pearson’s correlation with global FDR control at 0.05. There are 2103 CNA hubs in the marginal network with 47.9 percent of them having only one degree. In contrast, there are 15 CNA hubs in `prot-net` and only one of them has degree being one. The mean degree of the CNA hubs in the marginal network is 4, while for `prot-net` it is 39. Thus `spaceMap` is more likely to be able to narrow down the region of potential cancer drivers. This is due to `spaceMap` targeting direct interactions instead of marginal correlations, as well as utilizing group selection techniques. Moreover, by the module analysis described in Section 4.4, `prot-net` yielded 45 significant GO terms compared to 9 for the marginal network and thus is more functionally enriched. These observations imply that `spaceMap` is a useful addition to conventional pairwise correlation based analysis in integrating proteomics and CNA profiles and pinpointing important disease-relevant regulatory relationships.

Finally, among the CNA-protein edges in `prot-net`, the CNA profiles are positively correlated with their cis-regulated protein levels; whereas for trans-regulation, we did not observe a significant preference towards either positive regulation or negative regulation.

We also fit `scggm` to the CNA and protein levels of the BCPLS data set, but find evidence of high variability and instability. We first use 10-fold CV to choose `scggm`’s tuning parameters. However, this leads to a large number of protein–protein edges and very few CNA–protein edges ( first column of Table S.6). This is consistent with the observations from the simulation results of Section 2.1 and is likely due to high FDR in edge detection and lack of power in CNA-hub detection. In order to facilitate comparison with `spaceMap`’s `prot-net`, we instead choose `scggm`’s tuning parameters such that the resulting `Boot.Vote` network would have a similar size as `prot-net`. This leads to a network with 967 protein–protein edges, 574 CNA–protein edges (third column of Table S.6). There are many more edges if CV.Vote instead of `Boot.Vote` had been applied under these same tuning parameters (second column of Table S.6). This again indicates instability of the inferred network topology due to excess variability and overfitting. On the contrary, network topology inferred by `spaceMap` is reasonably stable: The number of protein–protein edges and CNA–protein edges are 1147 and 772, respectively, under CV.Vote; and 954 and 585 under `Boot.Vote`. We analyze `scggm`’s `Boot.Vote` network in more detail in Section S.3.2.

#### 2.2.2. RNA-net vs. prot-net

The protein and mRNA expressions are not jointly modeled due to limited sample size. Instead, we apply `spaceMap` to learn a separate RNA network, referred to as RNA-net. To facilitate comparison, `spaceMap` learned `RNA-net` through `Boot.Vote` (*B* = 1000) in such a way to produce similar edge sizes as `prot-net`; this resulted in 1010 RNA–RNA edges, 622 CNA–RNA edges, and 14 CNA-hubs. The list of the top-ranked CNA-hubs are shown in Table 2 and the complete list of the 14 CNA-hubs are provided in Table S.9. Module analysis reveals 13 modules (of size 15 or more) in `RNA-net`, with their enriched GO terms annotated in Table S.10.

`RNA-net` and `prot-net` share 6 common CNA-hubs (see boldfaced lines from Tables S.4 and S.9). Both identify the 17q12 amplicon as a major hub with shared cis regulatory elements such as ERBB2, GRB7, and PNMT. The common hub 5q34 has the largest out-degree in both networks. Other common CNA are 10p15.1-15.3, 15q13.1-15.1, 16q22.1-22.2, and 8q21.2-22.1. The CNA-hubs do not share many common targets, which is expected since there are less than 16% of common nodes (i.e., nodes corresponding to the same genes) in these two networks. There are 33 overlapping edges in total: 20 of them are CNA–expression edges, and 13 are expression–expression edges.

`prot-net` is more functionally-enriched compared to `RNA-net`, having 11 out of 15 CNA-hubs belonging to a module with at least one enriched GO term compared to only 2 out 17 for `RNA-net`. In addition, 45 GO terms are significantly enriched in `prot-net` modules, whereas only 24 enriched GO terms are detected in `RNA-net` modules. GO terms enriched in both networks include (innate) immune response, collagen catabolism, and extra-cellular matrix (ECM) organization. The last two are interesting as abnormal collagen bers in the ECM are known to play roles in invasive breast tumor activity (Grossman et al., 2016). The *GO-neighbor percentages* of CNA-hub neighborhoods also tend to be higher for `prot-net` (mean 63.25%) than `RNA-net` (mean 42.25%), as evidenced in Figure S.1.

### 2.3. OCPLS application

From Table 1, it can be seen that, compared with the breast cancer networks, the ovarian networks tend to have smaller CNA hubs. This is consistent with the marginal correlation based results from Mertins et al. (2016) and Zhang et al. (2016) for breast cancer and ovarian cancer, respectively. Moreover, the `prot-net` of ovarian cancer has 71 significant GO terms, whereas the `RNA-net` has none (Table S.12). This confirms the observation from the breast cancer application that the protein network tends to be more functionally enriched than the RNA network.

The network modules corresponding to the two leading CNA hubs — 20q11.22-.23 and 8q24.23-24.3 — in the ovarian CNA-protein network are illustrated in Figure S.7. Interestingly, two GO terms that are enriched in the two leading breast cancer CNA-hub modules, namely, RNA polymerase AI promoter and ion transmembrane transport, are also enriched in these two ovarian cancer CNA-hub modules (compare Figures 3 and S.7). The differences are, in breast cancer network, RNA polymerase II promoter genes are regulated by CNA in 5q34; and ion transmembrane transport genes are regulated by CNA in 17q21. These results suggest that crucial tumor related biological processes are triggered by different genetic alternation mechanisms in different types of cancers.

Among the four genes cis-regulated by the CNA-hub in 20q11.22-.23, RPRB1B, a novel gene also called CREPT and K-h, belongs to the RNA polymerase II promoter GO category. Specifically, RPRB1B increases cyclin D1 transcription during tumorigenesis, through enhancing the recruitment of RNAPII to the promoter region as well as chromatin looping (Lu et al., 2012). Another recent study further suggests the crucial role of RPRB1B in promoting repair of DNA double strand breaks and the potential of using RPRB1B as a biomarker to facilitate patient-specific individualized therapies (Patidar et al., 2016). Indeed, the specificity of the monoclonal antibody against CREPT has been recently characterized for preparation of industrial production (Ren et al., 2014). The fact that we successfully pin-point this important protein through our network analysis convincingly demonstrates the utility of `spaceMap` in revealing relevant and important biological information.

The other leading CNA-hub sits in 8q24.23-24.3, which is the only genome region that have shown frequent focal chromosome copy number gains in all four female cancer types, including ovarian, breast, endometrial and cervical cancers, in a very recent Pan-Cancer study (Kaveh et al., 2016). However, the functional consequences of copy number gain of this region remain largely unclear. Could the result of `spaceMap` network give us insights of the biological role of this important CNA region? Indeed, in the inferred ovarian CNA-protein network, we identified 11 cis-regulated proteins of this CNA-hub. The fact that this CNA-hub has the largest number of cis-regulations among all CNA-hubs in both the breast and ovarian CNA-protein networks also implies the uniqueness of this CNA region. Among the 11 cis-regulated proteins, three —TSTA3, NAPRT, and CYC1 — are from the Metabolic pathway. Specifically, TSTA3 controls cell proliferation and invasion and has been reported to exert a proto-oncogenic effect during carcinogenesis in breast cancer (Sun et al., 2016). NAPRT has been observed to promote energy status, protein synthesis and cell size in various cancer cells, and NAPRT-dependent NAD+ biosynthesis contributes to cell metabolism as well as DNA repair process, so that NAPRT has been recently suggested to be used to increase the efficacy of NAMPT inhibitors and chemotherapy (Piacente et al., 2017). CYC1 plays important roles in cell proliferation and comedo necrosis through elevating oxidative phosphorylation activities, and has been recently suggested to serve as a biomarker to predict poor prognosis in breast cancer patients (Han et al., 2016; Chishiki et al., 2017). All these are useful pieces of information to help to figure out the functional consequences of the amplification of the CNA-hub on 8q24.23-24.3, which then could lead to detection of novel drug targets or biomarkers for ovarian cancer.

## 3. Discussion

The proposed model `spaceMap` (Section 4.1) can successfully address many of the challenges inherent to learning networks from multiple -omic data types. Statistical efficiency is gained by discarding irrelevant interactions through a conditional graphical model framework. Moreover, pseudo-likelihood approximations and the MAP penalty increase power of CNA-hub detection compared with a likelihood-based CG model `scggm`. This has been convincingly shown by the simulation results in Section 2.1 as well as the BCPLS and OCPLS applications in Sections 2.2 and 2.3. Providing biological interpretation is another challenge after a network is learned. For this purpose, we develop a network analysis toolkit that facilitates the researchers to interpret their results and integrates with visualization software (Section 4.4).

We engineer the `spaceMap` R package to be computationally efficient. Model fitting steps are implemented with *Rcpp* and *RcppArmadillo* C++ bindings to R (Eddelbuettel, 2013; Eddelbuettel and Sanderson, 2014). Model selection procedures CV.Vote and `Boot.Vote` leverage R’s parallel processing backends. Detailed documentation and vignettes of the `spaceMap` R package and the network analysis toolkit are hosted on https://topherconley.github.io/spaceMap/. Details of the BCPLS application is illustrated in the *neta-bcpls* repository on GitHub (https://topherconley.github.io/neta-bcpls/).

In the applications of breast and ovarian cancers, protein level data results in more functionally enriched networks than RNA expression data for both cancer types, suggesting that protein data could be more informative for characterizing functional consequences of genetic alterations. While different sets of CNA-hubs and network modules are identified in the breast and ovarian CNA-protein networks, respectively, the leading modules in both networks are enriched with genes from the same GO categories, namely, ion transmembrane transport and regulation from RNA polymerase II promoter. This suggests a hypothesis that, although different cancers show large tumor heterogeneity in terms of genomic alterations, these distinct alteration events may eventually contribute to the same set of crucial biological processes. Therefore, identifying and characterizing the downstream crucial biological processes might be the key to design effective diagnosis and treatment strategies to overcome the challenges due to tumor heterogeneity in clinical practice. And network-based-system-learning approaches, like the one proposed in this paper, would be essential for such a goal.

Due to the small sample sizes, here we focus only on the most robust signals in the data and our top priority has been to control false discovery rate of the inferred networks. Although we are not expecting to capture all important regulatory patterns in such a way, we can be reasonably confident with those that are identified. In the future, with more proteomics samples available for cancer studies, we expect that `spaceMap` will be able to draw more insights on driver genes and genomic events in cancer.

Since `spaceMap` infers conditional dependency relationships, we expect that the interactions inferred by `spaceMap` are more direct than those inferred by marginal analysis. Specifically, compared to the results of marginal analysis using the same data by `spaceMap`, we find that `spaceMap` is more advantageous in narrowing down to small genome regions as tran-hub and thus could shed more lights on regulatory mechanisms of CNA on protein or RNA. However, this is at the cost of a more complicated model and consequently demands both more samples and computational efforts to fit the model.

In principle, `spaceMap` may also be used to conduct eQTL analysis using protein levels or RNA expressions (as response) and SNV (as predictor). Due to the large number of SNV, one may need to reduce the set of SNV through filtering and/or grouping.

## 4. Materials and Methods

### 4.1. spaceMap Model

Graphical models provide compact and visually intuitive representations of interactions (conditional dependency relationships) among a set of nodes (random variables). For example, CNAs and protein levels may be the nodes, while the edges may encode interactions among these molecular entities. Graphical models carry an advantage over relevance networks based on pair-wise correlations by being able to distinguish marginal dependency from conditional dependency. For instance, protein_*j*_ and protein_*k*_ may be marginally dependent because they are co-regulated by CNA_*l*_. But after conditioning on the effect of CNA_*l*_, they may become conditionally independent. Thus the interactions inferred by graphical models are more direct interactions com-pared with those based on marginal statistics such as pairwise correlations.

The goal is to learn the graph structure (ie. the set of edges) based on the observed data. In the following, we use 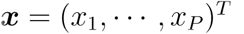 to denote one type of nodes, e.g., CNAs, and 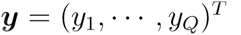 to denote another type of nodes, e.g., protein levels. Let 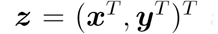 and assume that the random vector *z* has mean 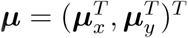 and a joint covariance matrix

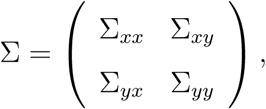

where *μ*_*x*_, *μ*_*y*_ are the mean vectors of *x* and *y*, respectively; ∑_*xx*_, ∑_*yy*_ are the covariance matrices of *x* and *y*, respectively; and 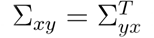is the covariance between *x* and *y*. Furthermore, the inverse of the joint *covariance matrix*, referred to as the concentration matrix, can be partitioned accordingly:

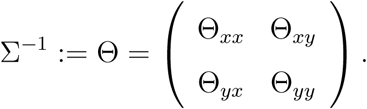

The off-diagonal entries of Θ are proportional to the *partial correlations*, i.e., the correlations between pairs of variables after removing linear effects of the rest of the variables. Under Gaussianity, partial correlations are the same as conditional correlations and thus zero entries in Θ mean that the respective pairs of random variables are conditionally independent given the rest of the variables. In contrast, a nonzero entry of Θ means conditional dependency and corresponds to an edge in the graph. Therefore, the goal of graph inference can be achieved by identifying nonzero entries of the concentration matrix Θ.

When there are two types of nodes, sometimes we are only interested in certain subsets of interactions. In the following, assume that we are only interested in the interactions among the *y* variables and those between the *y* variables and the *x* variables, but not the interactions among the *x* variables. Then there is no need to learn the entire concentration matrix Θ. Note that for 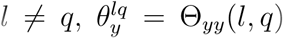 is proportional to 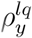 which denotes the partial correlation between *y*_*l*_ and *y*_*q*_ given 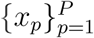 and 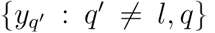; and in parallel, 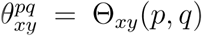 is proportional to 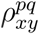 which denotes the partial correlation between *y*_*q*_ and *x*_*p*_ given 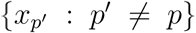 and 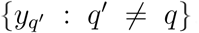. Therefore, we only need to elucidate the zero patterns of Θ_*yy*_ and Θ_*xy*_. Models for such a purpose are referred to as conditional graphical (CG) models.

For simplicity of notation, hereafter, assume that the mean vectors *μ*_*x*_ = 0, *μ*_*y*_ = 0. In Zhang and Kim (2014), it is assumed that given *x*, the conditional distribution of *y* is:

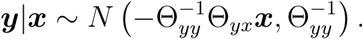

Parameters Θ_*yy*_ and Θ_*yx*_ are then learned by minimizing the negative conditional log-likelihood with *ℓ*_1_ penalties on Θ_*xy*_, Θ_*yy*_ to encourage sparsity. This method is called `scggm` (Sparse Conditional Gaussian Graphical Model).

In the following, we propose an alternative approach, called `spaceMap`, which uses regularized multivariate regression with sparsity- and hub- inducing penalties (Peng et al., 2010) to learn the zero patterns of Θ_*yy*_ and Θ_*xy*_. Unlike `scggm`, `spaceMap` does not rely on the conditional Gaussianity assumption for model fitting.

Peng et al. (2009) show partial correlations can be an efficient parameterization for graphical model learning. This is done through the connection between partial correlations and the regression coefficients while regressing each variable to the rest of the variables. This formulation can also be motivated through pseudo-likelihood approximations. Peng et al. (2009) further show this regression-based approach achieves higher power in edge detection compared to a likelihood-based approach when the true network exhibits hub structures, which is often the case for a GRN.

In `spaceMap`, we extend this regression framework to learn CG models. Note that, while regressing *y*_*q*_ to the rest of the nodes, we have:

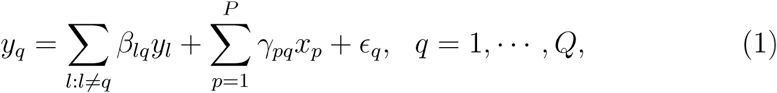

where the residual *∊*_*q*_ is uncorrelated with 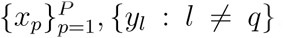, and the regression coefficients follow:

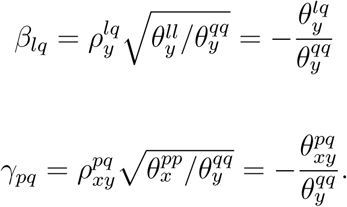

Since the diagonal entries 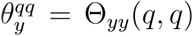 are positive, identifying nonzero entries of Θ_*yy*_ and Θ_*xy*_ is equivalent to identifying nonzero regression coefficients in (1). Also note that, equation (1) holds without the Gaussianity assumption.

We propose to minimize the following penalized *ℓ*_2_ loss criterion with respect to 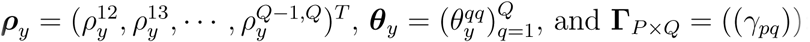:

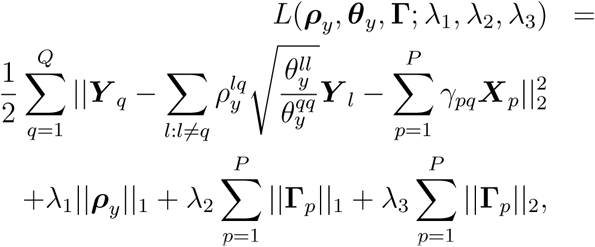

where 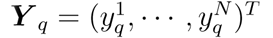and 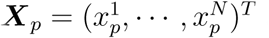 are observed samples of the nodes *y*_*q*_ and *x*_*p*_, respectively; Γ_*p*_ denotes the *p*th row of 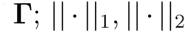 denote *ℓ*_1_ and *ℓ*_2_ norms, respectively; and λ_1_ > 0, λ_3_ > 0, λ_3_ > 0 are tuning parameters.

The *ℓ*_1_ norm penalty on *ρ*_y_ encourages sparsity in *y* — *y* interactions: When λ_1_ is sufficiently large, only some pairs of *y* nodes (but not all) will have the corresponding partial correlations estimated to be nonzero, thus having an edge in the inferred *y* — *y* network. By imposing regularization on the partial correlations *ρ*_*y*_ instead of on the *y* — *y* regression coefficients *β*_*lq*_’s, we not only reduce the number of parameters by nearly a half, but also ensure the *sign consistency*, due to 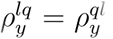 Moreover, since 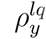 are on the same scale, the amount of regularization is comparable for different pairs of *y* nodes.

The combination of *ℓ*_1_ norm and *ℓ*_2_ norm penalties imposed on the *x* — *y* regression coefficients Γ encourages both sparsity in *x* — *y* interactions as well the detection of *x*-hubs: The *l*_1_ penalty induces overall sparsity of *x* nodes influencing the *y* nodes, while the *l*_2_ penalty encourages the selection of *x* nodes that have connections with many *y* nodes (i.e, *x*-hubs).

This model is referred to as `spaceMap` as it may be viewed as a hybrid of the space model (Peng et al., 2009) and the `remMap` model (Peng et al., 2010). Specifically, when λ_2_ = λ_3_ = 0, `spaceMap` reduces to a partial space model where only the *y* — *y* and *x* — *y* interactions are being fitted (but not the *x* — *x* interactions). On the other hand, the penalties on Γ is the same as the `MAP` penalty used in the `remMap` model which encourages the selection of *x*-hubs. `spaceMap` can be fitted through a *coordinate descent algorithm* similarly as in space and `remMap` by alternatively updating *ρ*_*y*_, *θ*_*y*_ and Γ.

For tuning parameters selection, we use K-fold cross validation (CV) to choose the optimal tuning parameters and adopt a sequential search strategy to efficiently navigate the 3D tuning grid. More details are given in section S.1 of the Supplementary Material. We then apply the `CV.Vote` procedure proposed in Peng et al. (2010) where only edges appearing in a majority of the cross validation networks are retained. The purpose of `CV.Vote` is to reduce the number of false positive edges. In application to real data, however, the false discovery rate (FDR) could still be high even after `CV.Vote` due to low signal-to-noise ratios and complicated noise structure. Therefore, we also consider a bootstrap-based aggregation procedure where we fit a network on each of the *B* bootstrap resamples of the data. We then only retain edges appearing in at least half of these networks (i.e., using a majority voting rule). We refer to this procedure as `Boot.Vote`. A similar strategy has been studied in Li et al. (2013) and is shown to be effective in reducing FDR. Compared to `CV.Vote`, as shown by the simulation results, `Boot.Vote` better controls FDR at the expense of computation and a small cost in power.

### 4.2. Simulation

We conduct two simulation studies to examine how well `spaceMap`, space, and `scggm` perform at network inference. In each simulation, one hundred independent replicates with sample size *N* = 250 are generated from a zero-mean multivariate Gaussian distribution. Each method is applied to each replicate to infer a network, where model tuning parameters are selected through 10-fold cross validation. The final network inference uses aggregation techniques `CV.Vote` and `Boot.Vote` to control the false discovery rate (FDR); The inferred networks are then compared with the true network to obtain power and FDR in terms of edge detection across all replicates. Moreover, *Matthew’s correlation coefficient (MCC)* is calculated to summarize the confusion matrix (Baldi et al., 2000); or in other words, the power-FDR tradeoff. MCC ranges from -1 to 1, where 1 corresponds to perfect match and −1 corresponds to perfect mismatch.

In the first simulation, referred to as `hub-net`, we generate a network with *P* = 35 CNA nodes, *Q* = 485 protein nodes and 577 total edges, following the hub network simulation from Peng et al. (2009) (with small modifications). Among the CNA nodes, 15 of them have at least 11 edges (CNA-hubs), and the rest has no edge (background nodes). The protein–protein network consists of five disjoint modules with around 100 nodes in each module since many real-world large networks exhibit modular structures. Moreover, the node degree follows a power-law distribution. The non-zero partial correlations fall in (−0.67,−0.1] ∪ [0.1, 0.67), with two modes around −0.28 and 0.28, respectively. Details on how the partial correlations are generated can be found in the Supplementary Material.

In the second simulation, referred to as `power-net`, we generate a network with more complicated structure: It has about 10 times as many non-hub CNA nodes and each module has roughly double the number of edges. This renders a power law network with the power parameter approximately 2.6. Specifically, there are 700 nodes (*P* = 210 CNA nodes and *Q* = 490 protein nodes) and 1257 edges. There are 10 CNA-hubs perturbing a subset of the 490 proteins. Among the 200 non-hub CNA nodes, 28 are confounders, meaning that they are correlated with a CNA-hub; 172 are background nodes, meaning that they are *not* connected to the rest of the CNA nodes and the protein nodes, although the background nodes are correlated among themselves. Further details of network generation and construction of the corresponding precision matrix Θ are given in Supplementary Section S.2.

### 4.3. Application: breast cancer and ovarian cancer proteogenomics

We apply `spaceMap` to the BCPLS data (Mertins et al., 2016) and the OCPLS data (Zhang et al., 2016) to demonstrate `spaceMap` ’s ability in identifying major functional consequences of DNA copy number alterations in a sample-size limited and biologically-heterogeneous context.

Protein abundance levels of 77 breast cancer tumor samples were obtained from the supplementary table of (Mertins et al., 2016). Protein levels of 174 ovarian cancer tumor samples were obtained from the supplementary table 2, sheet 2, of (Zhang et al., 2016). The corresponding level-three RNA-seq and segmented DNA copy number profiles were downloaded from TCGA web site (http://tcga-data.nci.nih.gov/tcga/). Global normalization were then performed to both the protein level and gene expression data sets.

Due to the relatively small sample sizes, genome-wide network reconstruction is not advisable. Instead, we focus on the most robust signals afforded in the data. This is accomplished by filtering out features with high missing rate and then selecting the most highly variable features, resulting in 1595 proteins and 1657 RNA expressions for breast cancer and 2097 proteins and 2236 RNAs for ovarian cancer, respectively. We also clustered CNAs into 1662 and 1349 larger genomic segments for breast cancer and ovarian cancer, respectively, using the fixed order clustering method proposed in Wang (2010), which helps to reduce multicollinearity among CNA features due to physical proximity (see section S.8).

### 4.4. Network interpretation toolkit

To facilitate biological interpretation, we built a toolkit to derive biological insights from the inferred networks. The application of the toolkit is illustrated under ’Network Analysis’ of Figure 1. The toolkit utilizes R package *igraph* (Csardi and Nepusz, 2006) to map user-supplied annotations onto the network and perform a rich suite of network analysis options. If the annotation contains gene coordinates, the toolkit also identifies cis/trans regulatory information. In our analysis, we define cis regulation to be within a ±4Mb window. In literature, there is not a standard window size for defining cis regulation. A large range of window sizes varying from 100k to 10Mb has been used in the past (Blackburn et al., 2015). Here we chose 4Mb as a trade-off between an overly liberal cis window and the risk of missing a cis regulation. The toolkit can conduct two types of analysis, one based on hubs and another based on modules.

In the hub analysis, we define any CNA node with at least one edge to an expression node (protein or RNA) as a *CNA-hub* and its corresponding *CNA-hub neighborhood* consists of all expression nodes that are directly connected to the CNA-hub by an edge. The toolkit prioritizes CNA-hubs based on a stability metric by calculating the average degree rank across the networks built on bootstrap resamples of the data (by the `Boot.Vote` procedure). Higher priority CNA-hubs have higher average rank, meaning that they have consistently high degree across the network ensemble. Next, the toolkit reports the number of cis/trans regulations found in each CNA-hub neighborhood and the number of protein/RNA nodes in cis with this CNA-hub from the entire network (referred to as *potential # of cis regulations*); see Table 2.

The hub analysis is enhanced if the annotation includes a functional mapping from databases like Gene Ontology (GO) to expression nodes. In our analysis, we construct a GO universe where each GO biological process term is required to have at least 15 participating genes in the network analysis, but no more than 300; for breast cancer, there are 167 and 129 biological processes meeting this criteria for protein and RNA, respectively; while for ovarian cancer there were 184 (protein) and 193 (RNA) biological processes. The toolkit reports the functional enrichment of a CNA-hub neighborhood through its *GO-neighbor percentage*: the percent of expression nodes in the neighborhood that share a common GO term with at least one other expression node neighbor; see Figure S.2.

Regarding module analysis, the toolkit evaluates which GO terms are significantly enriched in pre-specified network modules. In our analysis of the BCPLS data, we detect modules based on edge-betweenness (Newman and Girvan, 2004) using *igraph* — but other module-finding algorithms can be used. Modules with at least 15 nodes are subjected to GO enrichment testing through a hypergeometric test. GO terms are required to have at least 5 members in a module to be enriched. The FDR of GO enrichment is controlled at 0.05 (Benjamini and Hochberg, 1995). significantly-enriched modules, as well as any CNA-hub members of the modules, are organized into an accessible report as in Table S.5.

Taken together, the toolkit enables fast network analysis with well-organized results that is easily exported to other tools like *Cytoscape* (Shannon *et al.*, 2003), which rendered Figure 3.

## 5. Acknowledgments

Data used in this publication were generated by the National Cancer Institute Clinical Proteomic Tumor Analysis Consortium (CPTAC). This work has been supported by the Floyd and Mary Schwall Fellowship in Medical Research and grants NIH R01-GM082802, R01-GM108711, R01-CA189532 and NSF DMS-1148643. This work was also partly supported by grant U24 CA 210093, from the National Cancer Institute Clinical Proteomic Tumor Analysis Consortium (CPTAC).

## References

Akavia, U. D., Litvin, O., Kim, J., Sanchez-Garcia, F., Kotliar, D., Causton, H. C., Pochanard, P., Mozes, E., Garraway, L. A., Pe’er, D., 12 2010. An integrated approach to uncover drivers of cancer. Cell 143 (6), 1005–1017. URL http://www.sciencedirect.com/science/article/pii/S0092867410012936.

Baldi, P., Brunak, S., Chauvin, Y., Andersen, C. A. F., Nielsen, H., 2000. Assessing the accuracy of prediction algorithms for classification: an overview. Bioinformatics 16 (5), 412. URL http://dx.doi.org/10.1093/bioinformatics/16.5.412.

Barwe, S. P., Anilkumar, G., Moon, S. Y., Zheng, Y., Whitelegge, J. P., Rajasekaran, S. A., Rajasekaran, A. K., 2005. Novel role for na, k-atpase in phosphatidylinositol 3-kinase signaling and suppression of cell motility. Molecular biology of the cell 16 (3), 1082–1094.

Benjamini, Y., Hochberg, Y., 1995. Controlling the false discovery rate: A practical and powerful approach to multiple testing. Journal of the Royal Statistical Society. Series B (Methodological) 57 (1), 289–300. URL http://www.jstor.org/stable/2346101.

Blackburn, A., Almeida, M., Dean, A., Curran, J. E., Johnson, M. P., Moses, E. K., Abraham, L. J., Carless, M. A., Dyer, T. D., Kumar, S., Almasy, L., Mahaney, M. C., Comuzzie, A., Williams-Blangero, S., Blangero, J., Lehman, D. M., Goring, H. H. H., 9 2015. Effects of copy number variable regions on local gene expression in white blood cells of mexican americans. Eur J Hum Genet 23 (9), 1229–1235, article. URL http://dx.doi.org/10.1038/ejhg.2014.280.

Butte, A. J., Tamayo, P., Slonim, D., Golub, T. R., Kohane, I. S., 2000. Discovering functional relationships between rna expression and chemotherapeutic susceptibility using relevance networks. Proceedings of the National Academy of Sciences 97 (22), 12182–12186. URL http://www.pnas.org/content/97/22/12182.abstract.

Cancer Genome Atlas Network, et al., 2012. Comprehensive molecular portraits of human breast tumours. Nature 490 (7418), 61–70.

Cheng, J., Levina, E., Wang, P., Zhu, J., 2014. Sparse ising models with covariates. Biometrics 70 (4).

Chishiki, M., Takagi, K., Sato, A., Miki, Y., Yamamoto, Y., Ebata, A., Shibahara, Y., Watanabe, M., Ishida, T., Sasano, H., Suzuki, T., 2017. Cytochrome c1 in ductal carcinoma in situ of breast associated with proliferation and comedo necrosis. Cancer Science 108 (7), 1510–1519. URL http://dx.doi.org/10.1111/cas.13251.

Danaher, P., Wang, P., Witten, D., 2014. The jont graphical lasso for inverse covariance estiamtion across multiple classes. Journal of the Royal Statistical Society, Series B 76 (2).

Dimova, I., 2015. Whole genome microarray analysis in invasive ductal breast cancer revealed the most significant changes a ect chromosomes 1, 8, 17 and 20. International Journal of Sciences 4 (2015–01), 8–17.

Ellis, M. J., Gillette, M., Carr, S. A., Paulovich, A. G., Smith, R. D., Rodland, K. K., Townsend, R. R., Kinsinger, C., Mesri, M., Rodriguez, H., et al., 2013. Connecting genomic alterations to cancer biology with proteomics: the nci clinical proteomic tumor analysis consortium. Cancer discovery 3 (10), 1108–1112.

Friedman, J., Hastie, T., Tibshirani, R., 2008. Sparse inverse covariance estimation with the graphical lasso. Biostatistics 9 (3), 432–441.

Greenman, C., Stephens, P., Smith, R., Dalgliesh, G. L., Hunter, C., et al., 3 2007. Patterns of somatic mutation in human cancer genomes. Nature 446 (7132), 153–158. URL http://dx.doi.org/10.1038/nature05610.

Grossman, M., Ben-Chetrit, N., Zhuravlev, A., Afik, R., Bassat, E., Solomonov, I., Yarden, Y., Sagi, I., 2016. Tumor cell invasion can be blocked by modulators of collagen fibril alignment that control assembly of the extracellular matrix. Cancer Research 76 (14), 4249–4258. URL http://cancerres.aacrjournals.org/content/76/14/4249.

Haas, M., Wang, H., Tian, J., Xie, Z., 2002. Src-mediated inter-receptor cross-talk between the na+/k+-atpase and the epidermal growth factor receptor relays the signal from ouabain to mitogen-activated protein kinases. Journal of Biological Chemistry 277 (21), 18694–18702.

Han, Y., Sun, S., Zhao, M., et al., 2016. Cyc1 predicts poor prognosis in patients with breast cancer. Disease Markers 2016. URL https://doi.org/10.1155/2016/3528064.

Kaveh, F., Baumbusch, L. O., Nebdal, D., Børresen-Dale, A.-L., Lingjærde, O. C., Edvardsen, H., Kristensen, V. N., Solvang, H. K., 11 2016. A systematic comparison of copy number alterations in four types of female cancer. BMC Cancer 16 (1), 913. URL https://doi.org/10.1186/s12885-016-2899-4.

Kininis, M., Isaacs, G. D., Core, L. J., Hah, N., Kraus, W. L., 2009. Postrecruitment regulation of rna polymerase ii directs rapid signaling responses at the promoters of estrogen target genes. Molecular and cellular biology 29 (5), 1123–1133.

Kpetemey, M., Chaudhary, P., Van Treuren, T., Vishwanatha, J. K., 2016. Mien1 drives breast tumor cell migration by regulating cytoskeletal-focal adhesion dynamics. Oncotarget.

Li, L., Liu, B., Zhang, X., Ye, L., 2015. The oncoprotein hbxip promotes migration of breast cancer cells via gcn5-mediated microtubule acetylation. Biochemical and biophysical research communications 458 (3), 720–725.

Li, S., Hsu, L., Peng, J., Wang, P., 2013. Bootstrap inference for network construction. Annals of Applied Statistics 7 (1).

Li, Z., Langhans, S. A., 2015. Transcriptional regulators of na, k-atpase subunits. Frontiers in cell and developmental biology 3.

Litan, A., Langhans, S. A., 2015. Cancer as a channelopathy: ion channels and pumps in tumor development and progression. Frontiers in cellular neuroscience 9, 86.

Lu, D., Wu, Y., Wang, Y., Ren, F., Wang, D., Su, F., Zhang, Y., Yang, X., Jin, G., Hao, X., He, D., Zhai, Y., Irwin, D., Hu, J., Sung, J., Yu, J., Jia, B., Chang, Z., 2012. Crept accelerates tumorigenesis by regulating the transcription of cell-cycle-related genes. Cancer Cell 21 (1), 92–104. URL http://www.sciencedirect.com/science/article/pii/S1535610811004788.

Maurizio, E., Winiewski, J. R., Ciani, Y., Amato, A., Arnoldo, L., Penzo, C., Pegoraro, S., Giancotti, V., Zambelli, A., Piazza, S., Manfioletti, G., Sgarra, R., 2016. Translating proteomic into functional data: An high mobility group a1 (hmga1) proteomic signature has prognostic value in breast cancer. Molecular & Cellular Proteomics 15 (1), 109–123. URL http://www.mcponline.org/content/15/1/109.abstract.

Meinshausen, N., Bühlmann, P., 2006. High-dimensional graphs and variable selection with the lasso. The Annals of Statistics, 1436–1462.

Mertins, P., Mani, D. R., Ruggles, K. V., Gillette, M. A., Clauser, K. R., Wang, P., et al., 6 2016. Proteogenomics connects somatic mutations to signalling in breast cancer. Nature 534 (7605), 55–62, article. URL http://dx.doi.org/10.1038/nature18003.

Mijatovic, T., Ingrassia, L., Facchini, V., Kiss, R., 2008. Na+/k+-atpase subunits as new targets in anticancer therapy. Expert opinion on therapeutic targets 12 (11), 1403–1417.

Newman, M. E. J., Girvan, M., 2 2004. Finding and evaluating community structure in networks. Phys. Rev. E 69, 026113. URL http://link.aps.org/doi/10.1103/PhysRevE.69.026113.

Patidar, P. L., Motea, E. A., Fattah, F. J., Zhou, Y., Morales, J. C., Xie, Y., Garner, H. R., Boothman, D. A., 2016. The kub5-hera/rprd1b interactome: a novel role in preserving genetic stability by regulating dna mismatch repair. Nucleic Acids Research 44 (4), 1718–1731. URL http://dx.doi.org/10.1093/nar/gkv1492.

Paulovich, A. G., Billheimer, D., Ham, A.-J. L., Vega-Montoto, L., Rudnick, P. A., Tabb, D. L., Wang, P., Blackman, R. K., Bunk, D. M., Cardasis, H. L., et al., 2010. Interlaboratory study characterizing a yeast performance standard for benchmarking lc-ms platform performance. Molecular & Cellular Proteomics 9 (2), 242–254.

Peng, J., Wang, P., Zhou, N., Zhu, J., 2009. Partial correlation estimation by joint sparse regression models. Journal of the American Statistical Association 104 (486), 735–746, pMID: 19881892. URL http://dx.doi.org/10.1198/jasa.2009.0126.

Peng, J., Zhu, J., Bergamaschi, A., Han, W., Noh, D.-Y., Pollack, J. R., Wang, P., 03 2010. Regularized multivariate regression for identifying master predictors with application to integrative genomics study of breast cancer. Ann. Appl. Stat. 4 (1), 53–77. URL http://dx.doi.org/10.1214/09-AOAS271.

Piacente, F., Caffa, I., Ravera, S., Sociali, G., Passalacqua, M., Vellone, V. G., Becherini, P., Reverberi, D., Monacelli, F., Ballestrero, A., Odetti, P., Cagnetta, A., Cea, M., Nahimana, A., Duchosal, M., Bruzzone, S., Nencioni, A., 2017. Nicotinic acid phosphoribosyltransferase regulates cancer cell metabolism, susceptibility to nampt inhibitors, and dna repair. Cancer Research 77 (14), 3857–3869. URL http://cancerres.aacrjournals.org/content/77/14/3857.

Ren, F., Wang, R., Zhang, Y., Liu, C., Wang, Y., Hu, J., Zhang, L., Zhijie, C., 2014. Characterization of a monoclonal antibody against crept, a novel protein highly expressed in tumors. Monoclonal Antibodies in Immunodiagnosis and Immunotherapy 33, 401–408. URL https://doi.org/10.1089/mab.2014.0043.

Rothman, A., Levina, E., Zhu, J., 2010. Sparse multivariate regression with covariance estimation. Journal of Computational and Graphical Statistics 19 (4), 947–962.

Sahlberg, K. K., Hongisto, V., Edgren, H., Mäkelä, R., Hellstrom, K., Due, E. U., Vollan, H. K. M., Sahlberg, N., Wolf, M., Borresen-Dale, A.-L., et al., 2013. The her2 amplicon includes several genes required for the growth and survival of her2 positive breast cancer cells. Molecular oncology 7 (3), 392–401.

Samarakkody, A., Abbas, A., Scheidegger, A., Warns, J., Nnoli, O., Jokinen, B., Zarns, K., Kubat, B., Dhasarathy, A., Nechaev, S., 2015. Rna polymerase ii pausing can be retained or acquired during activation of genes involved in the epithelial to mesenchymal transition. Nucleic acids research 43 (8), 3938–3949.

Schäfer, J., Strimmer, K., 2004. An empirical bayes approach to inferring large-scale gene association networks. Bioinformatics 21 (6), 754. URL http://dx.doi.org/10.1093/bioinformatics/bti062.

Sun, Y., Liu, X., Zhang, Q., Mao, X., Feng, L., Su, P., Chen, H., Guo, Y., Jin, F., 4 2016. Oncogenic potential of tsta3 in breast cancer and its regulation by the tumor suppressors mir-125a-5p and mir-125b. Tumor Biology 37 (4), 4963–4972. URL https://doi.org/10.1007/s13277-015-4178-4.

Wang, P., 2010. Statistical Methods for CGH Array Analysis. VDM Verlag Dr.M IIer.

Wang, P., Chao, D., Hsu, L., 2011. Learning networks from high dimensional binary data: An application to genomic instability data. Biometrics 67 (1), 164–173.

Wol, A. C., Hammond, M. E. H., Hicks, D. G., Dowsett, M., McShane, L. M., Allison, K. H., Allred, D. C., Bartlett, J. M., Bilous, M., Fitzgibbons, P., et al., 2013. Recommendations for human epidermal growth factor receptor 2 testing in breast cancer: American society of clinical oncology/college of american pathologists clinical practice guideline update. Archives of Pathology and Laboratory Medicine 138 (2), 241–256.

Yuan, M., Lin, Y., 2006. Model selection and estimation in regression with grouped variables. Journal of the Royal Statistical Society: Series B (Statistical Methodology) 68 (1), 49–67.

Zhang, H., Liu, T., Zhang, Z., Payne, S. H., Zhang, B., McDermott, J. E., Zhou, J.-Y., Petyuk, V. A., Chen, L., Ray, D., Sun, S., Yang, F., Chen, L., Wang, J., Shah, P., Cha, S. W., Aiyetan, P., Woo, S., Tian, Y., Gritsenko, M. A., Clauss, T. R., Choi, C., Monroe, M. E., Thomas, S., Nie, S., Wu, C., Moore, R. J., Yu, K.-H., Tabb, D. L., Fenyo, D., Bafna, V., Wang, Y., Rodriguez, H., Boja, E. S., Hiltke, T., Rivers, R. C., Sokoll, L., Zhu, H., Shih, I.-M., Cope, L., Pandey, A., Zhang, B., Snyder, M. P., Levine, D. A., Smith, R. D., Chan, D. W., Rodland, K. D., 2016. Integrated proteogenomic characterization of human high-grade serous ovarian cancer. Cell 166 (3), 755–765. URL https://doi.org/10.1016/j.cell.2016.05.069.

Zhang, L., Kim, S., 02 2014. Learning gene networks under snp perturbations using eqtl datasets. PLoS Comput Biol 10 (2), e1003420. URL http://dx.doi.org/10.1371%2Fjournal.pcbi.100342.

